# Risk of hypoglycemia induced by pivalate-conjugated antibiotics in young children: a population-based retrospective study in Japan

**DOI:** 10.1101/350595

**Authors:** Tatebe Yasuhisa, Toshihiro Koyama, Naoko Mikami, Yoshihisa Kitamura, Toshiaki Sendo, Shiro Hinotsu

**Affiliations:** Department of Medicinal Pharmacology, Graduate School of Medicine, Dentistry and Pharmaceutical Sciences, Okayama University, Okayama, Japan; Department of Pharmacy, Okayama University Hospital, Okayama, Japan; Department of Pharmaceuticals Biomedicine, Graduate School of Medicine, Dentistry, and Pharmaceutical Sciences, Okayama University, Education and Research Center for Clinical Pharmacy, Graduate School of Medicine, Dentistry and Pharmaceutical Sciences, Okayama University, Okayama, Japan; Division of Pharmacy, Chiba University Hospital, Chiba, Japan; Center for Innovative Clinical Medicine, Okayama University Hospital, Okayama, Japan

**Keywords:** pivalate-conjugated antibiotics, carnitine deficiency, hypoglycemia, children, claims data

## Abstract

Infection is a common cause for an outpatient visit for young children. Pivalate-conjugated antibiotics (PCAs) are often used for these patients in Japan. However, a few case reports have shown that PCAs can provoke hypoglycemia in children, but no larger study has shown that PCAs increase the risk of hypoglycemia. The current study was performed as a retrospective review of children aged 1 month to 5 years old with at least once prescription of PCAs or other beta-lactam antibiotics from January 2011 to December 2013, using a medical and pharmacy claims database. Hypoglycemia was defined based on the International Statistical Classification of Diseases and Related Health Problems 10th Revision code or prescription of 10% or 20% glucose injection, and the incidence of hypoglycemic events was investigated. Logistic regression analysis was performed to examine the risk of hypoglycemia with PCAs compared with control antibiotics. The study cohort contained 179,594 eligible patients (male: 52.2%, mean age: 3.2 years). The numbers of prescriptions were 454,153 and 417,287 for PCAs and control antibiotics, respectively. Multivariate analysis showed that PCAs were associated with hypoglycemia (adjusted OR 1.18, 95% CI 1.12 to 1.24, *P* < 0.01), and the risk of hypoglycemia was also significantly increased with use of PCAs for ≤7 days (adjusted OR 1.17, 95% CI 1.11 to 1.24, *P* < 0.01). These results suggest that prescription of PCAs to young children should be avoided, even for a short time period.

## Introduction

Hypoglycemia is a common endocrine emergency and a major concern in neonates, infants and children because severe hypoglycemia can lead to neurological dysfunction such as neurocognitive defects, memory deficits, aphasia and hemiparesis (1, 2). In childhood, hypoglycemia is induced by various causes, including inborn errors of metabolism, infection, growth hormonal insufficiency, and adrenal insufficiency, as well as medications such as insulin derivatives and some antiarrhythmic agents (3–5).

In Japan, pivalate-conjugated antibiotics (PCAs) are widely used for pediatric patients with bacterial infections such as acute respiratory tract infection (ARTI) (6). Pivalic acid is conjugated to oral drugs to improve absorption. After absorption, the pivalic acid is liberated from these drugs, conjugated with carnitine in the liver, and excreted as the conjugate in urine, with a corresponding reduction of the serum carnitine concentration (7). Since carnitine plays an essential role in fatty acid oxidation in mitochondria (8), the reduced serum carnitine level can decrease lipid utilization for ATP production and result in impairment of gluconeogenesis, thereby inducing hypoglycemia (9).

PCAs have been reported to cause hypoglycemia in children in some cases, with a possible consequence of encephalopathy (10). Therefore, the Pharmaceutical and Medical Devices Agency (PMDA) and the Japan Pediatric Society (JPS) have cautioned that PCAs can induce hypoglycemia and recommended reduced use of these medicines when possible. However, while there are a few case reports of hypoglycemia induced by PCAs (10–12), no study has shown that PCAs increase the incidence of hypoglycemia compared with other beta-lactam antibiotics. Given that PCAs are widely prescribed in Japan, it is important to evaluate the risk of hypoglycemia with these antibiotics. Therefore, the aim of this study was to examine the risk of hypoglycemia associated with PCAs compared with other oral beta-lactam antibiotics using claims data in Japan.

## Methods

### Study design and data source

A retrospective cohort study was performed using a medical and pharmacy claims database constructed by the Japan Medical Data Center (JMDC, Tokyo, Japan). The Japanese healthcare system uses employee- or community-based plans with free access to hospitals and clinics. The JMDC database was launched in 2005 and includes multiple employee-based insurance plans, but not community-based plans. The population in the database had reached approximately 3 million at 2013, including a large pediatric population. The database contains data with an anonymized personal identifier, year of birth, gender, medical procedures, drugs prescribed and tests ordered, diagnosis, and diagnostic codes using the International Statistical Classification of Diseases and Related Health Problems 10th Revision (ICD10) (13).

### Drugs

Prescription data were used for PCAs, including cefcapene pivoxil, cefditoren pivoxil, cefteram pivoxil, and tebipenem pivoxil. All PCAs are classified as oral beta-lactam antibiotics and infections could be a risk factor of hypoglycemia. Therefore, we defined a control group of several beta-lactam antibiotics with no pivoxil residue, including amoxicillin, cefdinir, cefaclor, cefalexin, cefotiam, cefpodoxime proxetil, cefuroxime axetil, faropenem, and sultamicillin.

### Patients

Data from January 2011 to December 2013 were analyzed retrospectively. The subjects were children aged 1 month to 5 years old who had a prescription history of the beta-lactamantibiotics listed above. Neonates and patients with a prescription history of valproic acid, levocarnitine, or insulin derivatives in the period of diagnosis of hypoglycemia, or the diagnostic codes E10 (insulin-dependent diabetes mellitus), E70, E71, E72 (amino acids or fatty acid metabolic disorders), and E74 (abnormal carbohydrate metabolism), were excluded because valproic acid and levocarnitine affect the serum carnitine level, insulin frequently induces hypoglycemia, and metabolic diseases can change glucose metabolism.

### Definition of hypoglycemia

Hypoglycemia was defined as prescription of 10% or 20% glucose injection or the first record of ICD10 codes E160, E161 or E162, with exclusion of events if the diagnosis had no obvious association with PCAs; for example, hyperinsulinemia (E160). While hypoglycemia defined by diagnostic codes may reflect events of hypoglycemia in the real world to some extent (14), the identification of hypoglycemic episodes may not be sufficient because it is not necessary to mention a diagnosis of hypoglycemia in the medical receipt for a health insurance claim. In Japan, oral glucose or intravenous 10% or 20% glucose are typically administered to young children with symptomatic hypoglycemia. Thus, we also defined hypoglycemia in this study as intravenous glucose administration to elevate the power of detection. We did not define oral glucose intake as indicating a hypoglycemic event because it was difficult to identify oral supplementation with glucose using the JMDC database.

### Analysis

The incidence of defined hypoglycemia was examined within the study period. Eligible patients were divided into two groups based on prescription of PCAs or control antibiotics. Age; gender; number of days the drug was supplied; comorbidities causing hypoglycemia, such as hypopituitarism, adrenal insufficiency or type 2 diabetes mellitus; and conditions in which 10% or 20% glucose might be used, such as gastroenteritis and dehydration, were summarized using descriptive statistics. Each month in which a study drug was prescribed was counted as one exposure for each patient. If antibiotics in each group were concurrently prescribed for a patient in the same month, this exposure was excluded from data analysis. To exclude hypoglycemia possibly unrelated to antibiotics, antibiotic-related hypoglycemia was defined if the event occurred in the month of antibiotic exposure or in the following month.

Univariate analysis and multivariate logistic regression was performed to investigate factors associated with defined hypoglycemia. Covariates included gender, age, prescription of PCAs, and number of days of drug supply. To confirm that age and number of days of drug supply affected the risk of hypoglycemia induced by PCAs, multivariate analyses with stratification by age and number of days was performed. Adjusted odds ratios (OR), 95% confidence intervals (CI), and *P* values are reported. We also conducted univariate and subgroup analyses to examine the effects of confounding factors of gastroenteritis (ICD10 code: A0), dehydration (E86), adrenal insufficiency (E271, E272, E273, E274, E278, E279, E896), hypopituitarism (E230, E231, E233, E236, E237, E893) and type 2 diabetes mellitus (E10, E11, E12, E13, E14). For sensitivity analysis, hypoglycemia was redefined as diagnostic codes E160, E161, and E162 and intravenous 10% or 20% glucose prescriptions in the same month, and multivariate logistic analysis was conducted on these data as described above. All analyses were performed with JMP^®^ v.12 (SAS Institute Inc., Cary, NC, USA).

### Ethics statement

Data investigated in this study were deidentified by the JMDC in an unlinked manner, and therefore, informed consent was not required from patients. The study protocol and exemption of informed consent were approved by the Ethics Committee of Okayama University Graduate School of Medicine, Dentistry and Pharmaceutical Sciences and Okayama University Hospital.

## Results

### Patients

A total of 179,594 patients (male, 93,743 [52.2%]) were eligible for the study. The mean age (±SD) at dispensing was 3.16 (±1.55) years old (Table 1). A total of 328 patients were excluded from analyses due to a prescription history of insulin derivatives, valproate, or levocarnitine. The total number of visits with an antibiotics prescription were 871,440, including 454,153 for PCAs and 417,287 for control antibiotics. The mean age and proportion of males in the PCA group were significantly higher than those in the control group. The number of days of drug supply was lower in the PCA group, but the background of patients was slightly different in the two groups. Prescription patterns over time were similar in each group (Fig. 1).

**Table 1.** Characteristics of study patients in the Japan Medical Data Center claims database.

| Characteristics | Control group | PCA group |
| --- | --- | --- |
|  | n = 134,412 | n = 142,065 |
| Number of visits<sup>a</sup> | 417,267 | 454,153 |
| Age |  |  |
| Mean (SD) | 3.1 (1.55) | 3.21 (1.55) |
| Infant (%) | 33,569 (8.0) | 31,463 (6.9) |
| 1–2 years old (%) | 162,691 (39.0) | 169,619 (37.3) |
| 3–4 years old (%) | 158,426 (38.0) | 176,961 (39.0) |
| 5 years old (%) | 62,601 (15.0) | 76,110 (16.8) |
| Gender |  |  |
| Male (%) | 70,404 (52.4) | 74,606 (52.5) |
| Number of days of drug supply |  |  |
| Mean (SD) | 5.36 (3.9) | 4.90 (2.3) |
| ≤7 days (%) | 358,172 (85.83) | 409,182 (90.1) |
| 8–14 days (%) | 53,106 (12.73) | 42,328 (9.32) |
| ≥15 days (%) | 6,009 (1.44) | 2,643 (0.58) |
| Comorbidities |  |  |
| Dehydration (%) | 6,201 (1.49) | 8,680 (1.91) |
| Gastroenteritis (%) | 87,201 (20.9) | 89,593 (19.73) |
| Adrenal insufficiency (%) | 41 (0.01) | 61 (0.013) |
| Hypopituitarism | 194 (0.046) | 253 (0.056) |
| Type 2 diabetes mellitus (%) | 46 (0.011) | 47 (0.01) |

SD: standard deviation.
<sup>a</sup>Sum of visits by month that study drugs were prescribed.

**Figure 1.**
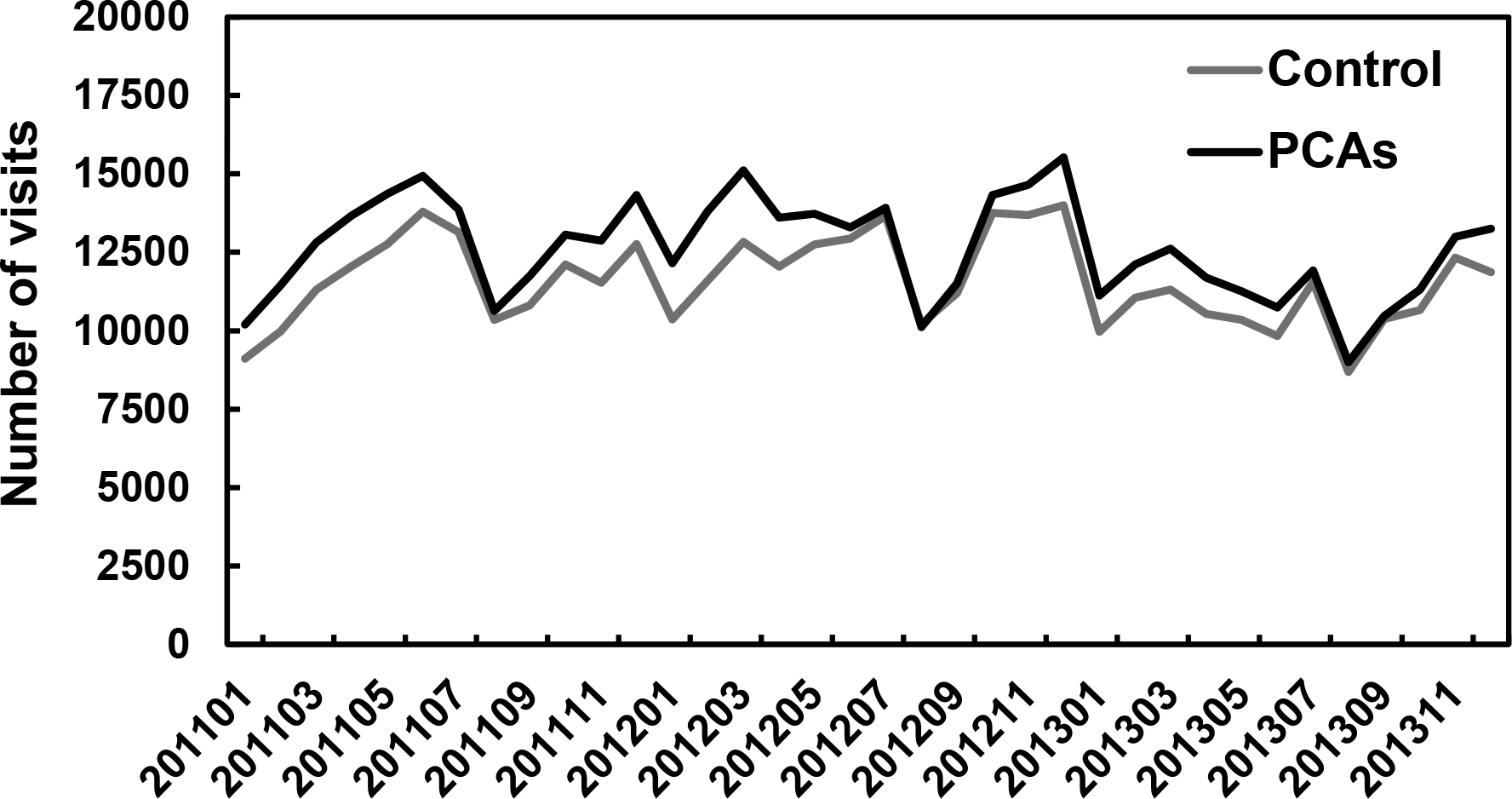
Prescription pattern of pivalate-conjugated antibiotics (PCAs) or control antibiotics in the study period. Black and grey lines represent the number of visits at which PCAs and control antibiotics were prescribed, respectively.

### Incidence of hypoglycemia

Univariate and multivariate logistic regression analyses revealed that the incidence of hypoglycemia (defined by ICD10 codes or prescription of 10% or 20% glucose injection) was higher in the PCA group compared with the control group (adjusted OR 1.18, 95% CI 1.12–1.24, *P* < 0.001, Table 2). In these analyses, male gender and age were associated with the risk of hypoglycemia (Table 2). While older age was related to a higher incidence of hypoglycemia, younger children, and especially infants, were more susceptible to the effect of PCAs on hypoglycemia (Table 3). The risk of hypoglycemia was also significantly increased with use of PCAs for ≤7 days, and this effect tended to increase for a longer period of drug supply (Table 3).

**Table 2.** Results of multivariate analysis for risk factors associated with hypoglycemia.

| Variable | Reference | OR (95% CI) | Adjusted OR<br>(95% CI) | *P* value |
| --- | --- | --- | --- | --- |
| Antibiotics |  |  |  |  |
| PCAs | Control | 1.19 (1.13–1.25) | 1.18 (1.12–1.24) | <0.001 |
| Gender |  |  |  |  |
| Male | Female | 1.14 (1.08–1.20) | 1.14 (1.08–1.20) | <0.001 |
| Age |  |  |  |  |
| 1–2 years old | Infant | 1.00 (0.90–1.12) | 1.00 (0.96–1.10) | 0.938 |
| 3–4 years old | Infant | 1.37 (1.23–1.53) | 1.36 (1.22–1.52) | <0.001 |
| 5 years old | Infant | 1.32 (1.18–1.49) | 1.31 (1.17–1.48) | <0.001 |
| Number of days drug supplied |  |  |  |  |
| 8–14 days | ≤7 days | 1.05 (0.97–1.13) | 1.06 (0.98–1.15) | 0.132 |
| ≥15 days | ≤7 days | 0.93 (0.71–1.20) | 0.96 (0.75–1.28) | 0.623 |

OR, odds ratio; CI, confidence interval; SD, standard deviation; PCAs, pivalate-conjugated antibiotics.

**Table 3.** Effect of PCAs on the risk of hypoglycemia with stratification by age or number of days of drug supply.

| Variable | Adjusted OR (95% CI) | *P* value |
| --- | --- | --- |
| Age |  |  |
| 0 years old | 1.32 (1.08–1.63) | 0.007 |
| 1–2 years old | 1.20 (1.10–1.32) | <0.001 |
| 3–4 years old | 1.15 (1.06–1.24) | <0.001 |
| 5 years old | 1.16 (1.03–1.31) | 0.017 |
| Number of days drug supplied |  |  |
| ≤7 days | 1.17 (1.11–1.24) | 0.007 |
| 8–14 days | 1.22 (1.05–1.41) | 0.011 |
| ≥15 days | 1.57 (0.90–2.69) | 0.11 |

OR, odds ratio; CI, confidence interval; PCAs, pivalate-conjugated antibiotics.

For sensitivity analysis, we redefined hypoglycemia as an event with a simultaneous ICD code and prescription of 10% or 20% glucose injection. The results of multivariate logistic regression analysis showed that the incidence of hypoglycemia with this definition was still higher in the PCA group (adjusted OR 1.29, 95% CI 1.11–1.49, *P* < 0.001, Table 4).

**Table 4.** Results of multivariate analysis as sensitivity analysis for hypoglycemia defined as prescription of 10 or 20% glucose injection and by ICD10 code.

| Variable | Reference | Adjusted OR<br>(95% CI) | *P* value |
| --- | --- | --- | --- |
| Antibiotics |  |  |  |
| PCAs | Control | 1.29 (1.11–1.49) | <0.001 |
| Gender |  |  |  |
| Male | Female | 1.26 (1.09–1.46) | 0.002 |
| Age |  |  |  |
| 1–2 years old | Infant | 1.35 (0.96–1.96) | 0.085 |
| 3–4 years old | Infant | 1.91 (1.37–2.76) | <0.001 |
| 5 years old | Infant | 1.72 (1.20–2.53) | 0.003 |
| Number of days drug supplied |  |  |  |
| 8–14 days | ≤7 days | 1.14 (0.91–1.41) | 0.258 |
| ≥15 days | ≤7 days | 1.18 (0.54–2.22) | 0.645 |

OR, odds ratio; CI, confidence interval; PCAs, pivalate-conjugated antibiotics

Patients with gastroenteritis and dehydration are often treated with glucose-containing fluids, and these events might have been included as antibiotics-induced hypoglycemia. Moreover, the hypoglycemia was caused by other factors (Table 5). Therefore, we conducted subgroup analyses in patients without dehydration, gastroenteritis, adrenal insufficiency, hypopituitarism, or type 2 diabetes mellitus. The incidence of hypoglycemia was still higher in the PCA group compared with the control group in all of these subgroup analyses (Table 6).

**Table 5.** Influence of comorbidities on the risk of hypoglycemia.

| Variable | OR (95% CI) | *P* value |
| --- | --- | --- |
| Dehydration | 33.6 (31.8–35.6) | <0.001 |
| Gastroenteritis | 3.13 (2.97–3.30) | <0.001 |
| Adrenal insufficiency | 4.40 (1.40–13.9) | 0.033 |
| Hypopituitarism | 2.65 (1.32–5.33) | 0.013 |
| Type 2 diabetes mellitus | 4.84 (1.53–15.3) | 0.026 |

OR, odds ratio; CI, confidence interval.

**Table 6.**
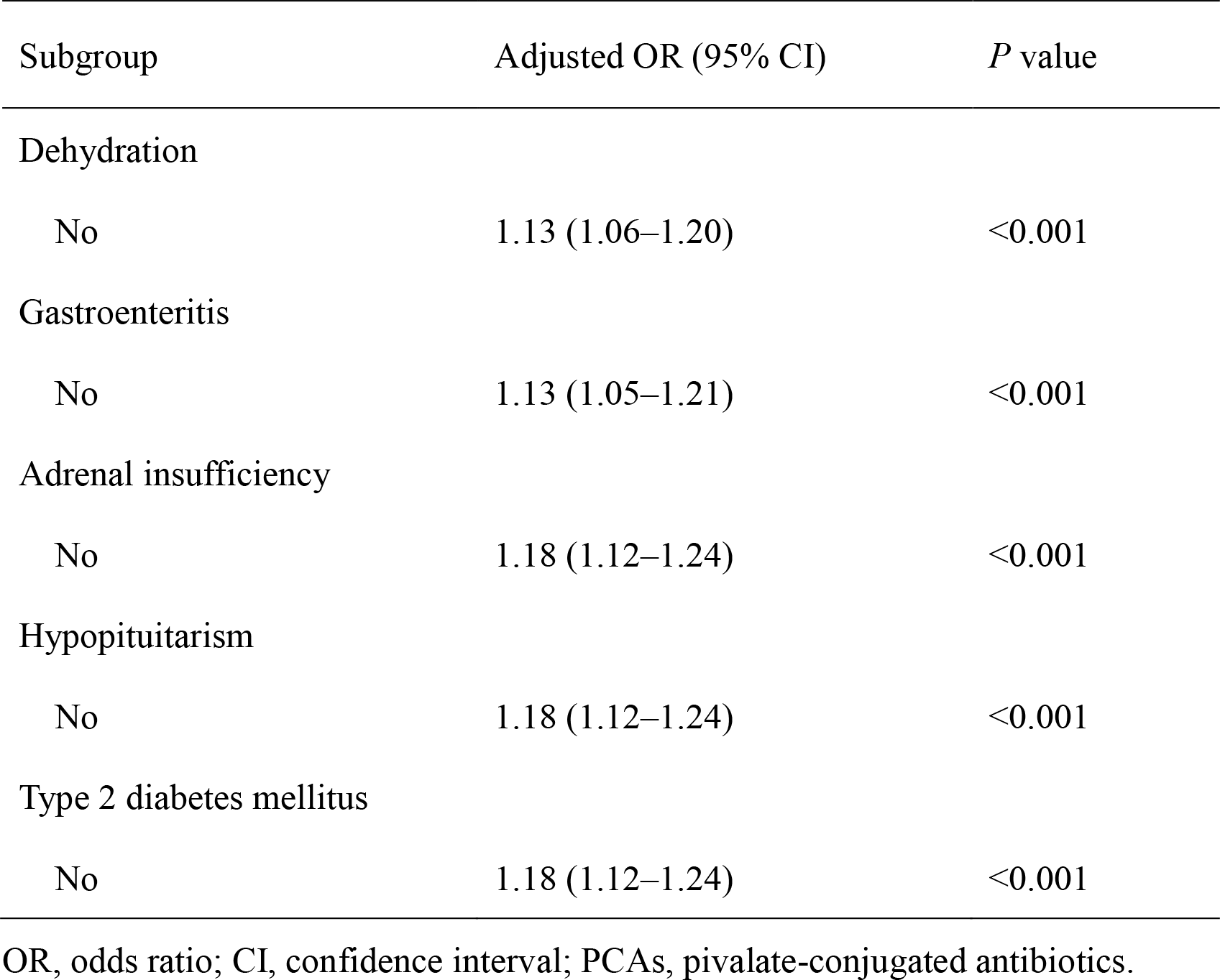
Risk of hypoglycemia induced by PCAs in subgroups of patients without comorbidities.

| Subgroup | Adjusted OR (95% CI) | *P* value |
| --- | --- | --- |
| Dehydration |  |  |
| No | 1.13 (1.06–1.20) | <0.001 |
| Gastroenteritis |  |  |
| No | 1.13 (1.05–1.21) | <0.001 |
| Adrenal insufficiency |  |  |
| No | 1.18 (1.12–1.24) | <0.001 |
| Hypopituitarism |  |  |
| No | 1.18 (1.12–1.24) | <0.001 |
| Type 2 diabetes mellitus |  |  |
| No | 1.18 (1.12–1.24) | <0.001 |

OR, odds ratio; CI, confidence interval; PCAs, pivalate-conjugated antibiotics.

## Discussion

This retrospective population-based study using the JMDC claims database showed that prescription of PCAs increased the incidence of hypoglycemia, defined as a prescription of 10% or 20% glucose injection or based on ICD10 codes, compared with other oral beta-lactam antibiotics in young children. These results were reproduced in subgroup and sensitivity analyses. In addition, we found that prescription patterns of PCAs and other antibiotics has not changed despite the warnings of the PMDA and JPS concerning PCA induced hypoglycemia since April 2012.

Several reports have shown that long exposure to pivalic acid can induce hypoglycemia following hypocarnitinemia (11, 12, 15), but in the current study we found that even use of PCAs for ≤7 days significantly increased the incidence of hypoglycemia (Table 3). Previous studies found that short term administration of pivampicillin and pivmecillinam resulted in a reduction of the mean serum creatine concentration to 15% of the pretreatment value in seven girls over a long period (16), and 7-day use of pivmecillinam reduced the serum carnitine by about 30% (17). This suggests that PCA-induced hypoglycemia can occur in short-term use.

Febrile illnesses are common in young children and the cause of fever is viral infections in many cases. Therefore, use of antibiotics is usually unnecessary (18), but antibiotics are often used in practice. In the ambulatory setting, >50% of children diagnosed with ARTIs received antibiotic prescriptions in the United States, while the estimated prevalence of ARTIs is <30% (19). A recent study using an administrative claims database in Japan showed that antimicrobial agents were prescribed for >60% of pre-school children in the cohort and that most prescribed antimicrobial agents were third-generation cephalosporins (20). Our data showed that >50% of prescribed beta-lactam antibiotics were PCAs, and this was consistent across the study period (Fig. 1). Therefore, PCAs are among the most prescribed antibiotics to young children in Japan, but these broad-spectrum antibiotics are probably unnecessary in many cases (21). Given that the unnecessary antibiotics were mainly third-generation cephalosporins, including PCAs, the risk of hypoglycemia induced by PCAs is not negligible, even though the incidence of hypoglycemia differed slightly between the PCA and control groups.

There are several limitations in this study. First, we did not consider the effects of concomitant medicines on hypoglycemia, except for insulin derivatives, valproic acid and levocarnitine. Indeed, hypoglycemia could also be induced by drugs such as quinolones, cibenzoline, pentamidine, and beta blockers (5). However, because most young children do not have morbidities treated with these drugs and most quinolones are contraindicated for children in Japan, it is unlikely that these drugs were prescribed in our subjects. Moreover, since it is unlikely that use of these drugs would affect the selection of antibiotics, the proportion of patients taking these drugs was presumably similar in the two groups. Second, the true incidence of hypoglycemia could not be determined because this study was conducted using the JMDC claims database and hypoglycemia was detected by ICD10 codes or prescription of 10% or 20% glucose injection, without using laboratory data for blood sugar levels. Furthermore, the time relationship between PCA use and hypoglycemia was unclear because we did not have the accurate dates for prescription of antibiotics and onset of hypoglycemia. Moreover, we had limited information from the claims database to adjust for differences in confounding factors between the groups, although subgroup analysis showed that prescription of PCAs was also a risk for hypoglycemia in children without several comorbidities. However, Japanese guidelines for pediatric ARTI recommend use of amoxicillin and third-generation cephalosporins such as cefditoren pivoxil (7, 22), and the selection of antibiotics is often dependent on physician preference in clinical practice. Therefore, the choice of antibiotics was unlikely to have been influenced by comorbidities that induce hypoglycemia. In fact, comorbidities of patients in the study were similar in each group (Table 1). Therefore, our finding that prescription of PCAs was associated with increased hypoglycemic events in young children is likely to be reliable. This is also supported by results showing that infants with lower serum carnitine and those with long-term antibiotic prescriptions were more susceptible to pivalate-induced hypoglycemia (Table 3).

## Conclusion

This retrospective cohort study showed that prescription of PCAs in young children increases the risk of hypoglycemia, compared with other antibiotics. Within the limitations of the study, this finding suggests that use of PCAs for young children should be avoided, even for a short time period.

## Acknowledgements

The study was supported by JMDC. No grant from a funding agency in the public, commercial, or not-for-profit sectors was received. The authors have no conflicts of interest to declare.

The authors’ contributions were as follows: conception and design of the study: Tatebe, Mikami, and Hinotsu; collection and assembly of data: Tatebe and Hinotsu; analysis and interpretation of data: Tatebe, Koyama, and Hinotsu; drafting of the article: Tatebe, Koyama, Kitamura, and Hinotsu; critical revision of the article for important intellectual content: Hinotsu, Kitamura, and Sendo; final approval of the article: All authors.

